# Assessment of the effects of transthyretin peptide inhibitors in *Drosophila* models of neuropathic ATTR

**DOI:** 10.1101/354555

**Authors:** Lorena Saelices, Malgorzata Pokrzywa, Katarzyna Pawelek, David S. Eisenberg

## Abstract

Transthyretin amyloidosis (ATTR) is a fatal disease caused by the systemic aggregation and deposition of transthyretin (TTR), a blood transporter that is mainly produced in the liver. TTR deposits are made of elongated amyloid fibrils that interfere with normal tissue function leading to organ failure. The current standard care for hereditary neuropathic ATTR is liver transplantation or stabilization of the native form of TTR by tafamidis. In our previous work, we explored an additional strategy to halt protein aggregation by capping pre-existing TTR fibrils with structure-based designed peptide inhibitors. Our best peptide inhibitor TabFH2 has shown to be effective at inhibiting not only TTR aggregation but also amyloid seeding driven by fibrils extracted from ATTR patients. Here we evaluate the effects of peptide inhibitors in two *Drosophila* models of neuropathic ATTR and compared their efficacy with diflunisal, a protein stabilizer currently used off-label for the treatment of ATTR. Our peptide inhibitor TabFH2 was found the most effective treatment, which resulted in motor improvement and the reduction of TTR deposition. Our *in vivo* study shows that inhibiting TTR deposition by peptide inhibitors may represent a therapeutic strategy for halting the progression of ATTR.

**SIGNIFICANCE STATEMENT:** Familial Amyloid Polyneuropathy (FAP) is a hereditary condition caused by the deposition of transthyretin (TTR) in nerves. Marked by progressive deficit and disability, FAP has no cure and limited therapeutic options. The replacement of the production source of mutant TTR by liver transplantation and the stabilization of native TTR by compounds, current lines of treatment, often fail to halt disease progression. Previously, we discovered that two segments of TTR drive amyloid deposition, and designed structure-based peptide inhibitors. Here we evaluate these peptide inhibitors in FAP models of *Drosophila*. The most efficient inhibitor resulted in an improvement of locomotor abilities and a reduction of TTR deposition. This study points to peptide inhibitors as a potential therapeutic strategy for FAP.

## Introduction

Transthyretin amyloidosis (ATTR) is caused by the deposition of amyloid fibrils of transthyretin (TTR), a transporter of thyroxine and retinol in the blood and cerebrospinal fluid (Costa et al., 1978; Sletten et al., 1980; Westermark et al., 1990). Both mutant and/or wild-type TTR are found in these deposits, which are made of elongated, resilient amyloid fibrils. Although TTR deposits in virtually every organ, ATTR patients commonly exhibit progressive cardiomyopathy and/or polyneuropathy that lead to disability and death (Galant et al., 2017). In hereditary ATTR, deposition and disease progression is accelerated by TTR mutations thereby resulting in earlier onsets (Hurshman Babbes et al., 2008). The most common form of hereditary amyloidosis, ATTR-V30M, is associated with familial amyloid polyneuropathy (FAP) that typically manifests in the third or forth decade of life, with or without cardiac involvement (Gertz et al., 2015).

The current standard of care for hereditary ATTR cases is liver transplantation. Because TTR is mainly synthesized in the liver, this procedure replaces the circulating mutated TTR with the more stable wild-type TTR (Benson, 2012). Liver transplantation has prolonged the life of many ATTR patients, with the most favorable prognosis for FAP-V30M cases (Benson, 2013). However, this drastic procedure is not recommended for cases of advanced age, longstanding disease, or with cardiac involvement, factors that have been associated with higher post-surgical mortality (Carvalho et al., 2015). In patients with a significant cardiac involvement, amyloid derived from wild-type TTR produced by the transplanted liver may continue to add to existing cardiac deposits, leading to accelerated deposition and heart failure (Liepnieks and Benson, 2007). Liver transplantation remains the only specific therapy for a limited number of ATTR patients in many countries.

More recent efforts are focused on the stabilization of the native form of TTR by small compounds as a method to inhibit protein aggregation (Miroy et al., 1996). The functional native form of TTR is a tetramer made of two homo-dimers that form a hydrophobic tunnel in which thyroxine binds as it is transported (Nilsson et al., 1975). The aggregation of transthyretin to form amyloid fibrils starts by the dissociation of the tetramer into monomers that unfold to expose amyloidogenic segments (Foss et al., 2005). Kelly and coworkers found that compounds bound to the thyroxine site stabilize the tetramer, thereby inhibiting protein aggregation (Miroy et al., 1996; Baures et al., 1998). Out of those studies, a stabilizing compound known as Tafamidis was discovered and is now approved in Europe, Japan, Mexico, Argentina and other countries for FAP patients (Bulawa et al., 2012). Some other compounds have shown promising results in the stabilization of TTR *in vitro* and were tested in clinical trials. This is the case for diflunisal, a non-steroidal anti-inflammatory drug that is currently being used off-label for the treatment of cardiac amyloidosis (Castano et al., 2012).

In two recent studies, we have developed and optimized peptide inhibitors that inhibit both transthyretin aggregation *in vitro* as well as amyloid seeding catalyzed by ATTR *ex-vivo* amyloid fibrils (Saelices et al., 2018; Saelices et al., 2015). We first discovered that there are two amyloidogenic segments of TTR that drive protein aggregation: β-strands F and H (Saelices et al., 2015). In that study, we determined the atomic structures of these two segments in their amyloid form, which allowed us to design specific peptide inhibitors of F and H β-strands self-association. Later we further optimized these inhibitors to inhibit amyloid seeding driven by ATTR *ex-vivo* amyloid fibrils (Saelices et al., 2018).

*Drosophila melanogaster* has recently emerged as a convenient model for human transthyretin deposition disorder (Pokrzywa et al., 2007; Berg et al., 2009; Pokrzywa et al., 2010; Andersson et al., 2013; Iakovleva et al., 2015). The overexpression of several familial and engineered amyloidogenic variants of human TTR in neurons results in TTR deposition in the brain, fat body and glia, atrophy of wings, locomotor impairment and shortened lifespan. In this manuscript, we evaluate the efficacy of our peptide inhibitors and the stabilizing compound diflunisal in two *Drosophila* models of ATTR. We found that the treatment of diseased flies with our optimized peptide inhibitor results in motor improvement and a reduction of TTR deposition.

## Materials and Methods

### ANTIBODIES

Antibodies used were rabbit anti-human transthyretin polyclonal antibody (DAKO, Agilent Technologies; 1:2,000) and horseradish peroxidase-conjugated goat anti-rabbit IgG antibody (DAKO, Agilent Technologies; 1:5,000).

### *DROSOPHILA* STOCKS

The formation of intracellular amyloid aggregates in thoracic adipose tissue and brain glia in ATTR models of the fruit fly results in an abnormal wing posture and motor defects (Pokrzywa et al., 2007; Pokrzywa et al., 2010; Iakovleva et al., 2015). Several ATTR models are available to be tested in flies; here, the focus was on flies carrying the TTR familial mutant V30M (Iakovleva et al., 2015) (abbreviated V30M), and the amyloidogenic mutant V14N/V16E (Pokrzywa et al., 2007) (abbreviated TTR-A). Transgenic lines were generated in the w1118 strain. Two transgenes for the human TTR gene UAS-TTRV30M and UAS-TTRV14N/V16E (abbreviated UAS-TTR-A) were expressed under control of pan-neuronal GAL4 driver (nSyb-GAL4) to drive expression in all types of post-mitotic neurons. Genotypes: w; +; UAS-TTRV30M/nSyb-GAL4 (Iakovleva et al., 2015), or w; +; UAS-TTRV14N/V16E/nSyb-GAL4 (Pokrzywa et al., 2007); wild-type Oregon R strain was obtained from Drosophila Bloomington Stock Center (BDSC #6361, Indiana University) and used as healthy controls in crosses with the nSyb-GAL4 driver line (w; +; +/nSyb-GAL4).

### FLY REARING AND DRUG FEEDING

Flies were kept at 60% humidity at 20 °C under a 12:12 hour light:dark cycle (8 a.m. to 8 p.m. daily) until fly eclosion and at 29 °C post-eclosion. This temperature shift was adopted to lower the expression of nSyb-GAL4 driver during development before adding the tested compounds. The crossings were reared in bottles containing standard *Drosophila* food (corn meal, corn syrup solids, yeast, water, and agar). Newly eclosed female flies (10 flies per vial) were transferred into 5 ml ventilated vials (75 × 13 mm, polystyrene tubes with archiving caps with filter, Sarstedt, Nümbrecht, Germany), containing low-melt fly food and tested compounds according to the formula developed by Markstein et al. for mixing drugs in low volumes (Markstein et al., 2014). Briefly, the food was prepared with distilled water containing 2% (w/v) autoclaved yeast, 7% (v/v) corn syrup liquids, and 1.5% (w/v) agarose (composed of 1 part standard agarose to 11 parts low-melt agarose). The food was mixed as a liquid with drugs at 37 °C. The resulting food and compound mixtures solidified at 30 °C into soft fly edible gels. All peptides were dissolved in 0.22 μm filtered water, first to 5 mM stock solution. These working solutions were further diluted in fly food prior to use to final concentration: 100 μM or 300 μM. Flies were fed compounds present in fly food after adult flies eclosure (developmental stages excluded) until death. Fresh food containing the compounds was changed every second or third day and the number of dead flies was recorded.

### ANALYSIS OF LOCOMOTOR ABILITIES OF TREATED FLIES

All females flies were pooled and randomized. Ten flies of each genotype or drug treatment were placed in plastic vials in replicates, gently tapped down to the bottom and allowed to climb. Several motor skills were analyzed in a climbing assay at various time points along with survival analysis. (i) Mean distance or mean length of trajectories corresponding to one fly in a vial in mm. (ii) Mean velocity of 10 flies moving in one vial in mm/s. (iii) Motion represents the percentage of flies being in movement in one vial. A fly is considered in movement when its velocity is equal to or higher than 2.5 mm/s. (iv) Maximum velocity of flies moving in one vial in mm/s. (v) Mean length of trajectory among all detected flies in one vial in mm. (vi) Total distance or the sum of all trajectory lengths in one vial in mm. Recordings of 10-second fly movements were acquired in duplicates and analyzed with FlyTracker hardware and software (Pokrzywa et al., 2017). Data analysis was performed in real time and again in the offline mode.

### ANALYSIS OF TTR DEPOSITION IN HEAD HOMOGENATES FROM TREATED FLIES

For every condition, 20 heads were homogenized in 100 μl Triton X-100 buffer (1%Triton X-100, 1x PBS, pH 7,6) containing a protease inhibitor cocktail (general use, Amresco) on ice. Samples were mixed gently and centrifuged for 20 minutes at 15,000 × g at 4 °C. This process was done twice and the supernatant was collected and saved as the soluble fraction (SF). The pellet was resuspended in 100 μ1 of 50 mM Tris pH 7,6, 4% SDS, gently vortexed and saved as the insoluble fraction (IF). Both SF and IF samples were boiled separately for 10 minutes. Samples were sonicated for 10 minutes and centrifuged at 15,000 × g at room temperature for 10 minutes. Both supernatants were collected and saved (SF-S1 and IF-S1, respectively). The pellet resultant from SF was resuspended in 100 μ1 of 50mM Tris pH 7,6, 4% SDS (SF-P1). The pellet resultant from IF was resuspended in 50 μ1 of 50 mM Tris-HCl pH 7,6, 175 mM NaCl, 5 mM EDTA, 5% SDS, 8 M urea, and vortexed for 1.5 hours (IF-P1). SF-S1 and SF-P1 were mixed and loaded together. IF-S1 and IF-P1 were mixed and loaded together. 4 x LDS sample buffer and DTT containing (10 x) Sample Reducing Agent (ThermoFisher Scientific) were added to the samples. Samples were boiled for 20 minutes before electrophoresis. Total protein concentration was estimated with Pierce™ BCA Protein Assay Kit - Reducing Agent Compatible (ThermoFisher Scientific). For each treatment 2.2 μg protein was resolved on NuPAGE^®^ Novex^®^ 4-12% Bis-Tris Protein Gels in MES SDS running buffer and electroblotted onto a nitrocellulose membrane using iBlot2 gel transfer device (ThermoFisher Scientific). All steps were performed according to the manufacturer. The primary antibodies used were rabbit polyclonal against human TTR 1:2,000 (DAKO). Detection was performed with Western Breeze Chromogenic kit for Rabbit Primary Antibodies. Transthyretin levels from four independent blots were quantified with Gel-Doc XR+ Imager and Image Lab 5.2 software (Bio-Rad). Recombinant TTR wild-type protein standard was used to calculate absolute levels of TTR immunodetected on blots.

### EXPERIMENTAL DESIGN AND STATISTICAL ANALYSIS

Statistical analysis of fly survival was performed with IBM SPSS Statistics 20 for Windows (IBM Corporation). Survival data were analyzed with the Kaplan-Meier method, and statistical comparisons were made with Log rank pair-wise comparisons. Additionally, Cox Regression analysis was performed to estimate hazard coefficient for each treatment. Statistical analysis of motor skills under various treatments was performed with Prism 7 for Mac (OriginLab). Statistical significance for locomotor effects was determined by nonparametric one-way ANOVA for multiple comparisons. Groups were compared in post hoc analysis with Holm-Sidak correction. All samples and animals were included in the analysis. The mean difference was considered to be statistically significant at 95% confidence interval. All quantitative experiments are presented as means ± S.E.M. of at least 6 independent experiments. No statistical methods were used to predetermine sample size before analysis. The experiments were randomized. Vials were coded and investigators were blinded to allocations both during readouts and outcome assessment.

## Results

### We tested TTR peptide inhibitors in two *Drosophila* models of ATTR

The expression and aggregation of human mutant TTR in the fruit fly results in the reduction of both lifespan and climbing ability (Pokrzywa et al., 2007; Pokrzywa et al., 2010; Iakovleva et al., 2015). To evaluate the efficacy of TabFH2 *in vivo*, we made use of two Drosophila models carrying the following genotypes: (i) TTRV30M (here referred as V30M flies), expressing the familial TTR mutant V30M (Iakovleva et al., 2015); and (ii) TTR-A, expressing an engineered TTR variant carrying the double mutation V14N/V16E (Pokrzywa et al., 2007). We selected these two strains for our evaluation because they display different phenotype intensities, being the effects of V30M TTR expression on both lifespan and climbing ability more pronounced. After eclosion, flies were fed with vehicle, TabFH1, or TabFH2 at two different concentrations as outlined in Fig. 1a. Oregon wild-type flies were included in the climbing assay as healthy controls. TabFH1 was included in the analysis to confirm optimization and rule out nonspecific effects of peptide-based treatment. Diflunisal was included in the assay for efficacy comparison. TTR-expressing vehicle-treated flies were used as negative controls. Survival analysis showed that neither TabFH1 nor TabFH2 resulted in a significant effect (Fig. 1b). However, diflunisal treatment resulted in a prolonged lifetime only for V30M flies, but not for TTR-A (Fig. 1b), suggesting that this effect may be unrelated to TTR deposition.

**Figure 1.**
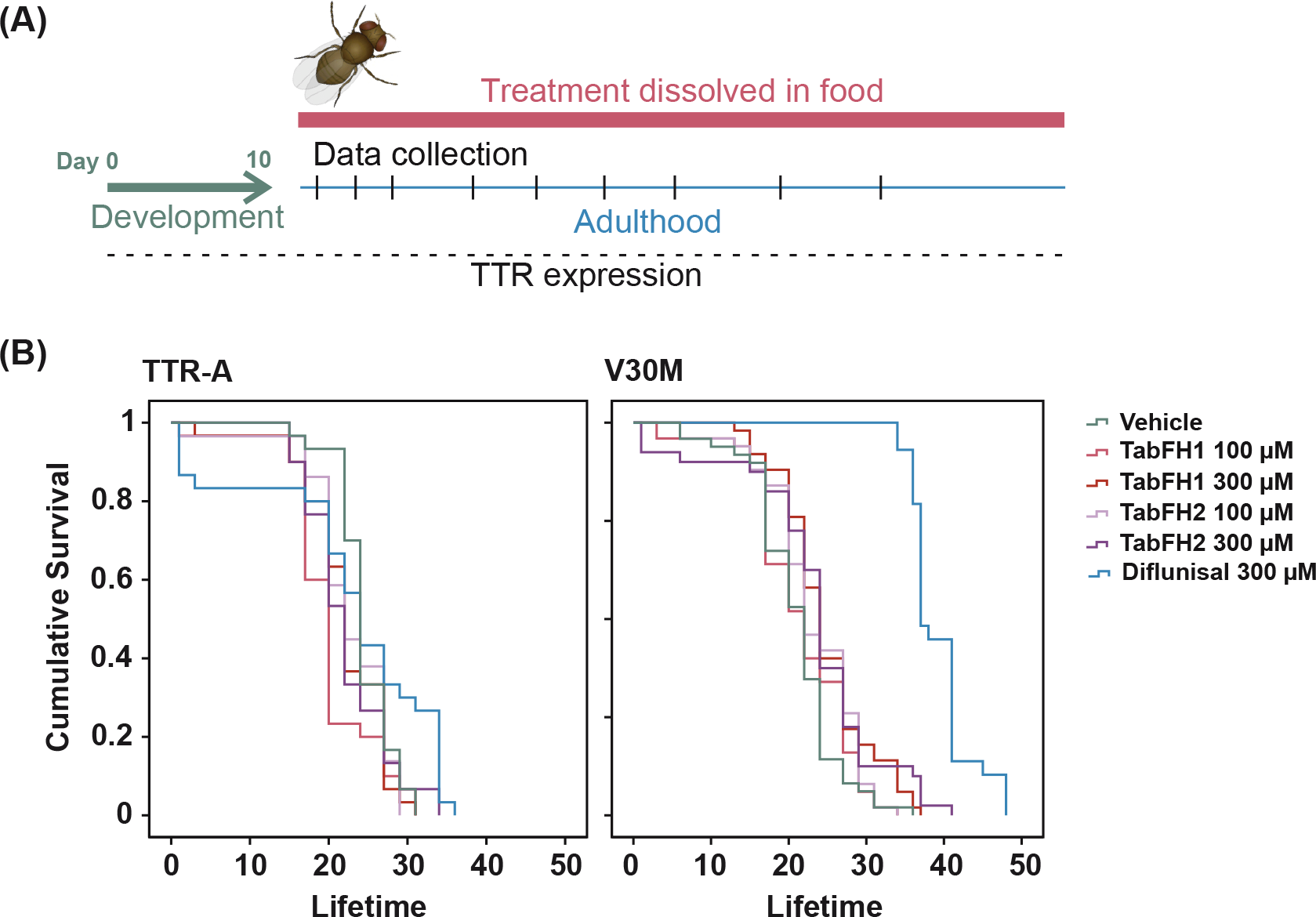
Treatment outline and lifespan evaluation of ATTR models of *Drosophila melanogaster*. **(A)** Treatment outline. Drugs were dissolved in fly food and flies were fed with the mix from day 1 after fly eclosion. Food was freshly made every two or three days. The flies were subjected to climbing assays and data were collected at different time points. **(B)** Lifespan analysis of TTR-A and V30M flies after treatment with vehicle, TabFH1, TabFH2 or diflunisal.

### The effect of TTR peptide inhibitors on fly mobility was evaluated in climbing assays at different time points

Ten flies of each genotype or drug treatment were placed at the bottom of plastic vials in replicates and allowed to climb, and several locomotor paradigms were assessed: mean distance or mean length of trajectories, mean velocity, overall motion that represents percentage of flies being in movement, maximum velocity, mean length of trajectory, and total distance or the sum of all trajectory lengths. 10-second recordings were acquired in duplicates and analyzed with FlyTracker hardware and software, as described before (Pokrzywa et al., 2017). Overall, TTR-A flies presented better climbing measures when compared to V30M flies (Fig. 2 vs. Fig. 3). For both genotypes, the computational analysis of climbing traces revealed that TabFH2 treatment results in a significant relative motor improvement in several parameters, including traveled distance, climbing velocity, and overall time of flies in motion (Fig. 2 and 3, Tables 1 and 2). The positive effect of diflunisal on climbing skills was significant, but lesser than TabFH2. As expected, treatment of flies with our peptide control TabFH1 showed a limited effect.

**Figure 2.**
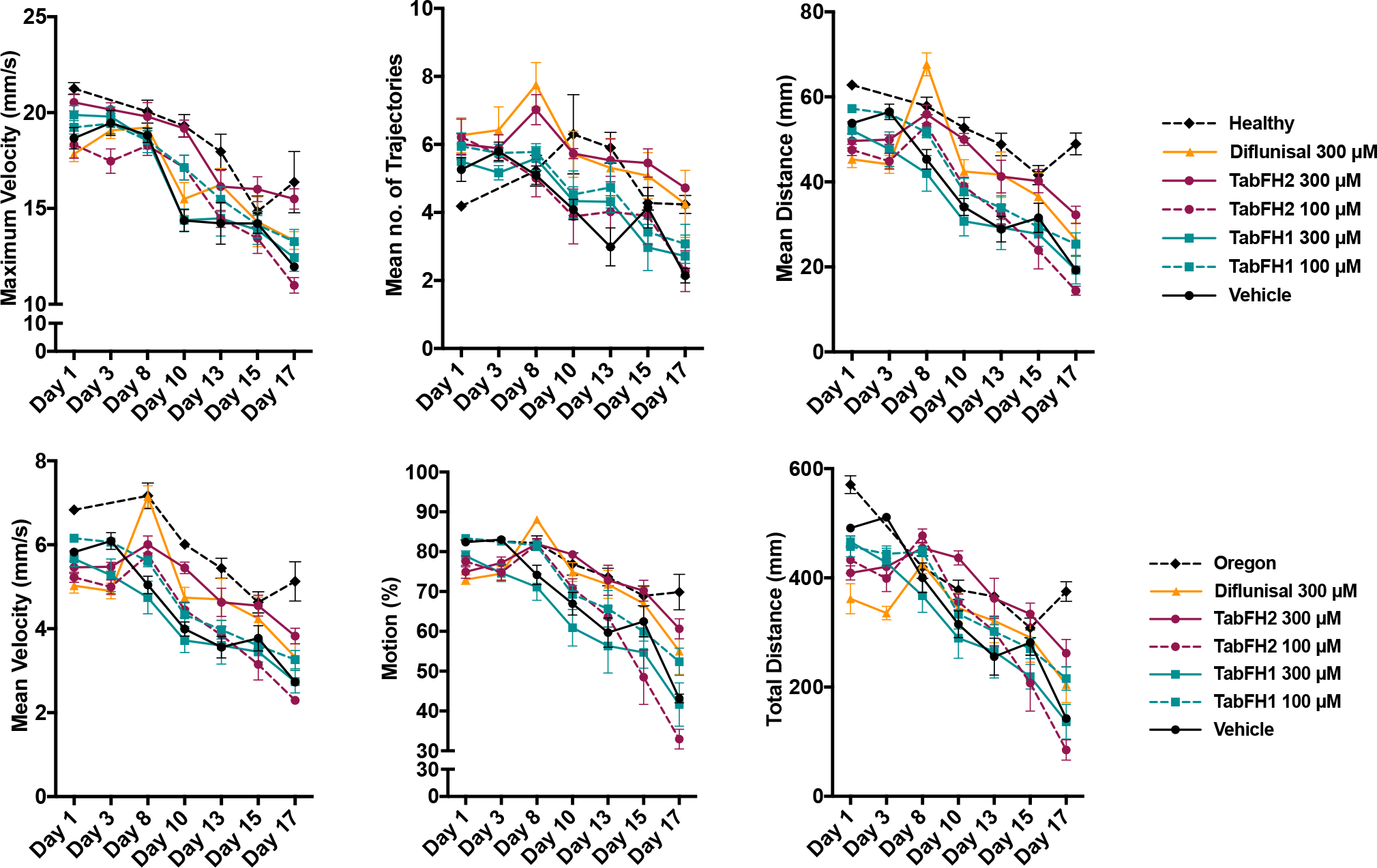
Effect of TabFH2 on TTR-A flies in climbing assays. Fly climbing abilities were assessed by measuring six motor parameters on Flytracker as described in Materials and Methods. Ten flies were gently tapped down in a vial in 6 replicates. The flies were subjected to climbing assays and data were collected at different time points. Dots represent the average of single means obtained from 6 independent measurements. Error bars, S.E.M.

**Figure 3.**
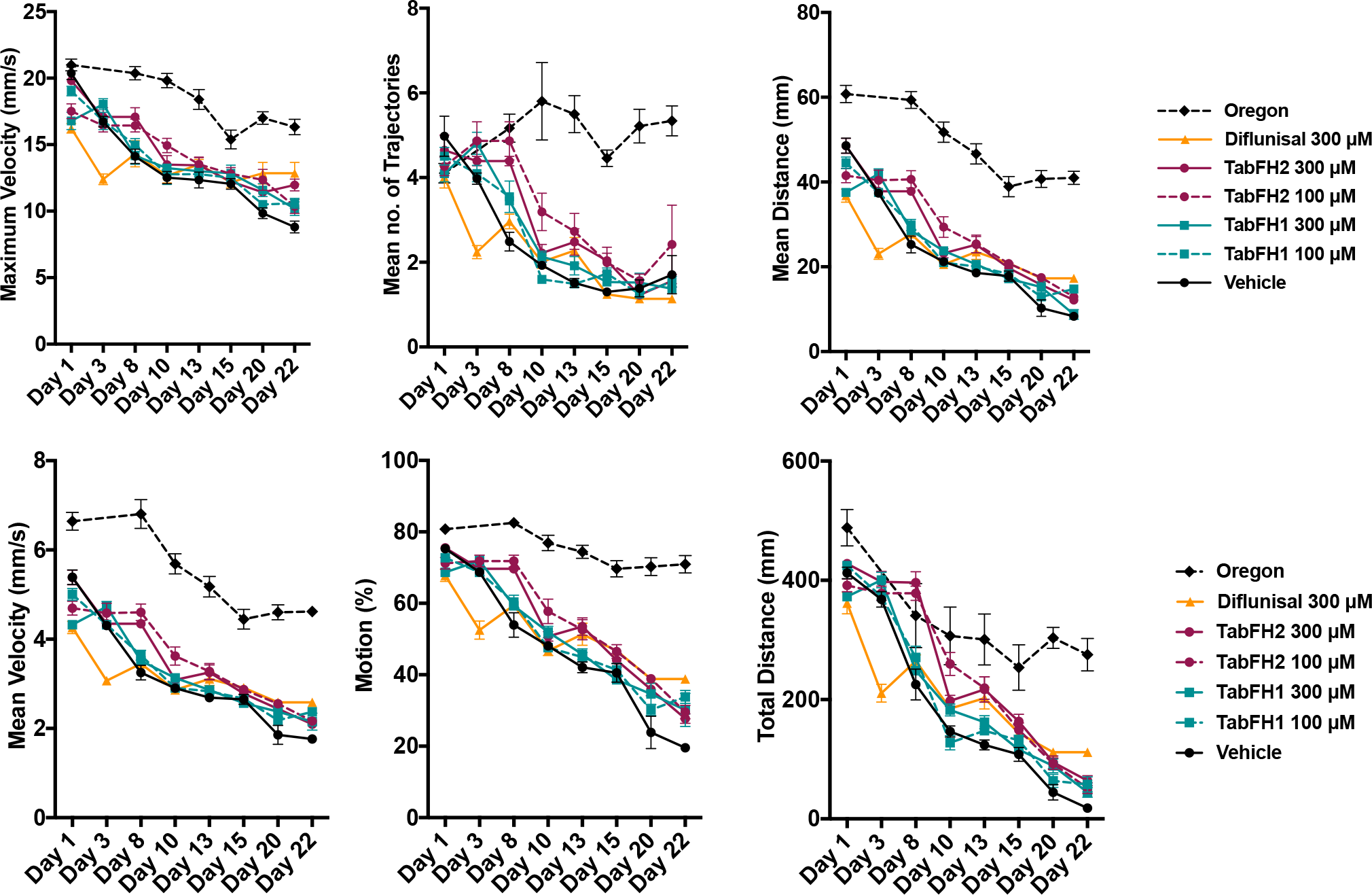
Effect of TabFH2 on V30M flies in climbing assays. Fly climbing abilities were assessed by measuring six motor parameters on Flytracker as described in Materials and Methods. Ten flies were gently tapped down in a vial in 8 replicates. The flies were subjected to climbing assays and data were collected at different time points. Dots represent the average of single means obtained from 8 independent measurements. Error bars, S.E.M.

**Table 1.**
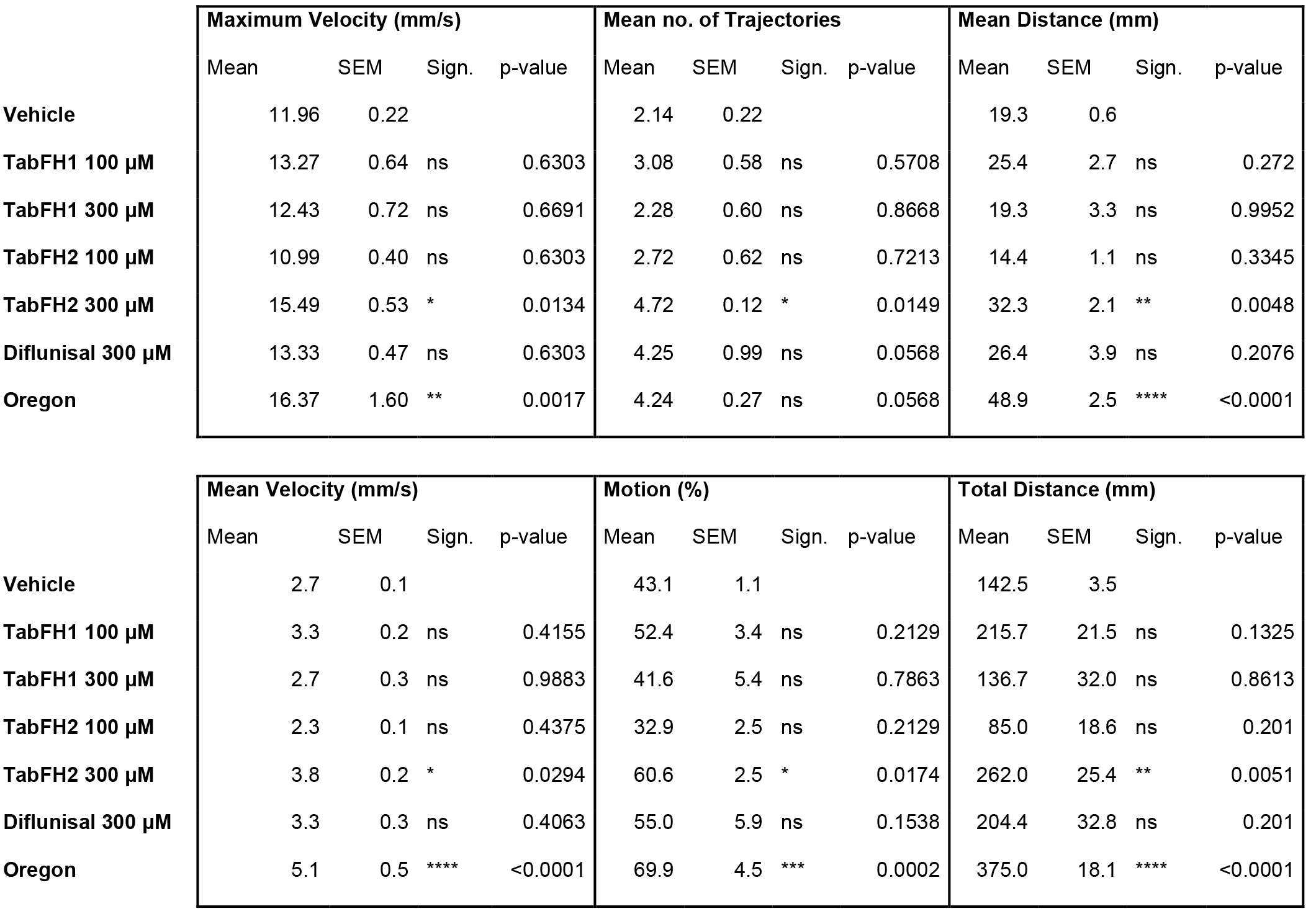
Statistical analysis of TTR-A flies after 17 days of treatment. Statistical significance for locomotor effects of each of the treatments was determined in comparison to vehicle-treated flies using nonparametric one-way ANOVA for multiple comparisons with Holm-Sidak correction. *p ≤ 0.05. **p ≤ 0.005. ***p ≤ 0.0005. ****p ≤ 0.0001.

**Table 2.** Statistical analysis of V30M flies after 22 days of treatment. Statistical significance for locomotor effects of each of the treatments was determined in comparison to vehicle-treated flies using nonparametric one-way ANOVA for multiple comparisons with Holm-Sidak correction. *p ≤ 0.05. **p ≤ 0.005. ***p ≤ 0.0005. ****p ≤ 0.0001.

|  | Maximum Velocity (mm/s) |  |  |  | Mean no. of Trajectories |  |  |  | Mean Distance (mm) |  |  |  |
| --- | --- | --- | --- | --- | --- | --- | --- | --- | --- | --- | --- | --- |
|  | Mean | SEM | Sign. | p-value | Mean | SEM | Sign. | p-value | Mean | SEM | Sign. | p-value |
| **Vehicle** | 8.81 | 0.44 |  |  | 1.71 | 0.45 |  |  | 8.32 | 0.66 |  |  |
| **TabFH1 100 µM** | 10.58 | 0.38 | * | 0.0486 | 1.54 | 0.13 | ns | 0.9551 | 14.77 | 0.97 | **** | <0.0001 |
| **TabFH1 300 µM** | 10.19 | 0.51 | ns | 0.0687 | 1.38 | 0.05 | ns | 0.9306 | 8.78 | 1.17 | ns | 0.7186 |
| **TabFH2 100 µM** | 10.36 | 0.52 | ns | 0.0687 | 2.42 | 0.93 | ns | 0.7528 | 12.92 | 0.77 | ** | 0.0022 |
| **TabFH2 300 µM** | 11.97 | 0.45 | *** | 0.0002 | 1.56 | 0.15 | ns | 0.9551 | 12.16 | 0.53 | ** | 0.0083 |
| **Diflunisal 300 µM** | 12.85 | 0.82 | **** | <0.0001 | 1.14 | 0.07 | ns | 0.8609 | 17.31 | 0.67 | **** | <0.0001 |
| **Oregon** | 16.33 | 0.59 | **** | <0.0001 | 5.34 | 0.35 | **** | <0.0001 | 40.98 | 1.58 | **** | <0.0001 |

**Table 2. Statistical analysis of V30M flies after 22 days of treatment.**
|  | Mean Velocity (mm/s) |  |  |  | Motion (%) |  |  |  | Total Distance (mm) |  |  |  |
| --- | --- | --- | --- | --- | --- | --- | --- | --- | --- | --- | --- | --- |
|  | Mean | SEM | Sign. | p-value | Mean | SEM | Sign. | p-value | Mean | SEM | Sign. | p-value |
| **Vehicle** | 1.76 | 0.05 |  |  | 19.6 | 1.3 |  |  | 18.0 | 4.8 |  |  |
| **TabFH1 100 µM** | 2.37 | 0.07 | *** | 0.0004 | 33.8 | 1.8 | *** | 0.0004 | 57.8 | 12.4 | * | 0.046 |
| **TabFH1 300 µM** | 2.15 | 0.19 | * | 0.0219 | 30.1 | 4.7 | ** | 0.0085 | 44.7 | 8.5 | ns | 0.0987 |
| **TabFH2 100 µM** | 2.17 | 0.09 | * | 0.0219 | 29.5 | 1.9 | ** | 0.0094 | 51.6 | 8.9 | ns | 0.0764 |
| **TabFH2 300 µM** | 2.10 | 0.05 | * | 0.0245 | 27.7 | 1.3 | * | 0.019 | 63.9 | 8.3 | * | 0.0226 |
| **Diflunisal 300 µM** | 2.59 | 0.06 | **** | <0.0001 | 38.8 | 1.2 | **** | <0.0001 | 111.7 | 4.0 | **** | <0.0001 |
| **Oregon** | 4.62 | 0.13 | **** | <0.0001 | 71.0 | 2.4 | **** | <0.0001 | 275.2 | 27.2 | **** | <0.0001 |

### The effects of TabFH2 on locomotor abilities were apparent after one week of treatment

2D drawings of climbing traces were generated by Flytracker software to better visualize the effect of the studied drugs on locomotor skills of diseased flies after 7 days of treatment. They show a prominent positive effect on both fly lines after treatment with TabFH2 (Fig. 4). This effect was more noticeable in TTR-A flies, whose behavior when treated with TabFH2 was comparable to healthy flies (Fig. 4A). In contrast, diflunisal treatment exhibits a significant positive effect in TTR-A flies after 7 days of treatment, but not in V30M flies (Fig. 4). Consistent with our locomotor assessment, climbing traces show that TabFH1 has a limited effect on treated flies (Fig. 4).

**Figure 4.**
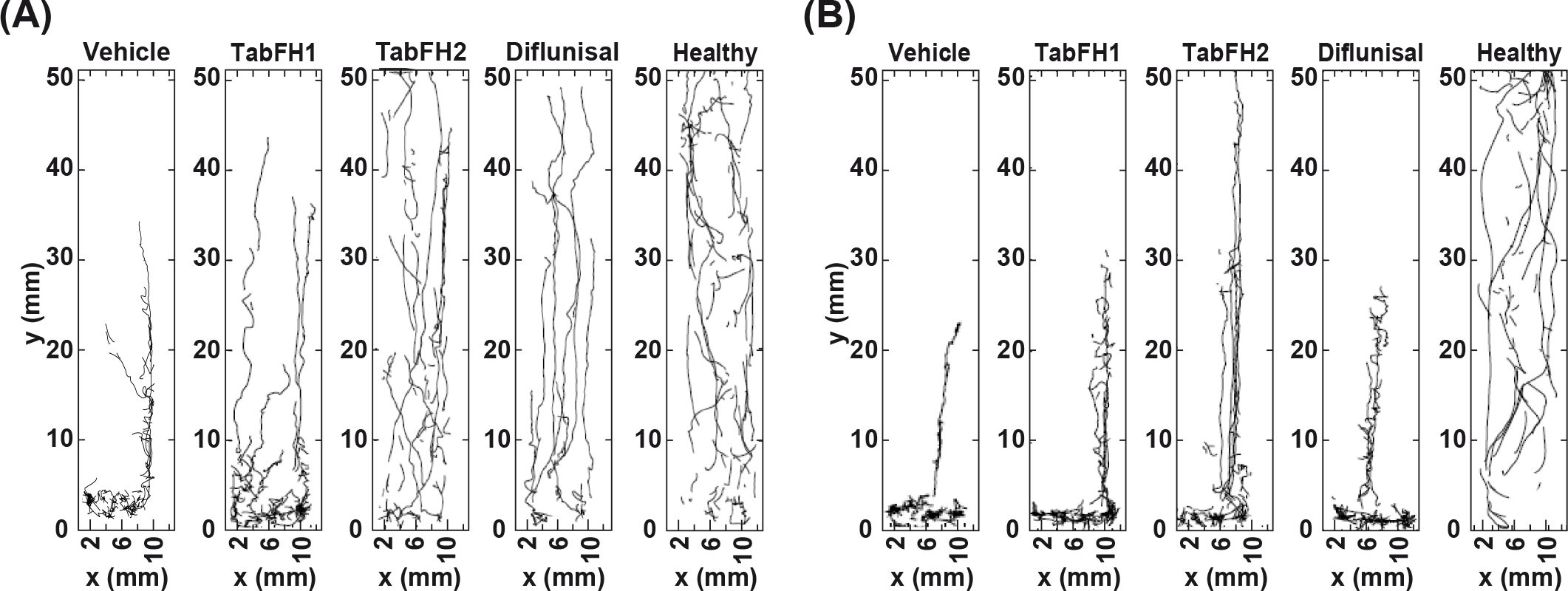
Two-dimensional drawings of climbing flies. 10-second trajectory traces were recorded from Oregon flies (labeled as healthy), TTR-A **(A)**, or V30M **(B)** flies after 7 days of treatment with vehicle or TabFH1, TabFH2, or Diflunisal at 300 μM.

### The improvement of locomotor skills was accompanied by a dose-dependent reduction of TTR deposition

For every condition, 20 heads were homogenized in a solution containing detergent and centrifuged twice to collect soluble and insoluble fractions. After further treatments of the samples to ensure denaturation, TTR deposition in the head of the treated flies was assessed by western blot (Fig. 5A,B). The results show a dose-dependent significant reduction of TTR insoluble deposition when flies are treated with our peptide inhibitors (Fig. 5C). Samples obtained from flies treated with diflunisal show a reduction of TTR insoluble fraction that is similar to that observed with TabFH2 at the same concentration (Fig. 5C). However, the overall expression of TTR in flies treated with diflunisal seemed to be lower than the other conditions (Fig. 5A,B).

**Figure 5.**
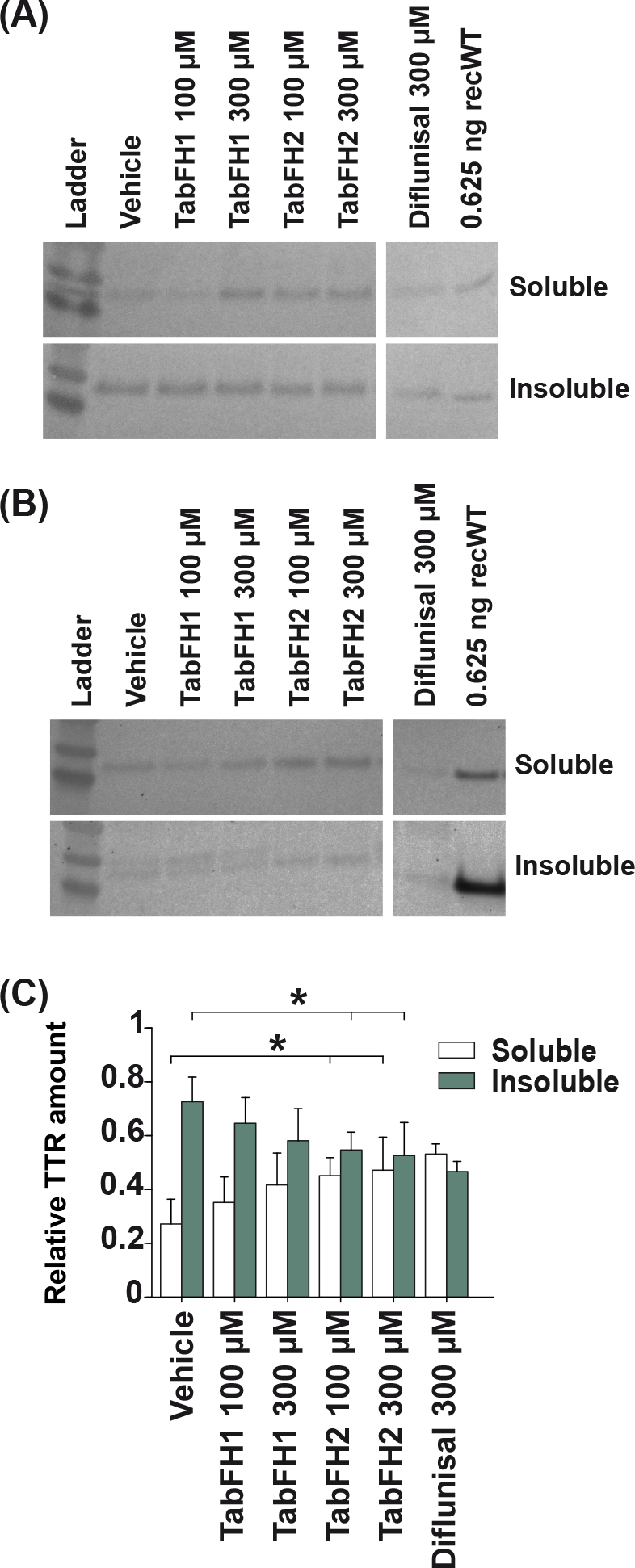
Evaluation of TTR deposition in the head of treated flies. Representative anti-TTR western-blots of soluble and insoluble material extracted from TTR-A **(A)** and V30M **(B)** fly heads. Every sample was extracted from 20 flies and 2.2 μg was loaded per condition. Anti-human transthyretin polyclonal antibody (DAKO) was used to detect transthyretin in the soluble and insoluble fractions. **(C)** Quantification of four independent western blots. Insoluble and soluble relative amounts were normalized to the total detected signal per condition. These fractions were obtained as described in Materials and Methods.

## Discussion

In this study, we evaluate peptide inhibitors of TTR aggregation in two *Drosophila* models of familial amyloid polyneuropathy. We found that TabFH2 not only reduces TTR deposition in treated flies but also results in the amelioration of the locomotor impairment observed when flies are untreated.

TabFH2 was found to be the most efficient drug in this study, followed by diflunisal and TabFH1 in this order. TabFH2 showed the greatest effect on delaying locomotor impairment and diminishing TTR deposition after continuous treatment (Fig 2, 3 and 5). Although the locomotor improvement in flies treated with diflunisal was significantly lower than with TabFH2, we did not observe any significant differences in TTR relative deposition between these two drugs (Fig. 5C). However, western blots show a lower overall presence of both soluble and insoluble TTR in flies treated with diflunisal vs. other treatments (Fig. 5A,B). Consistent with our previous *in vitro* work, the peptide control TabFH1 shows a limited capacity to inhibit TTR deposition (Fig. 5C) and no significant effect on the disease-related locomotor decline (Fig. 2 and 3) (Saelices et al., 2018).

The analysis of lifespan after treatment with peptide inhibitors and diflunisal resulted in a puzzling observation. Flies treated with the most efficient drug of the study, TabFH2, did not exhibit an elongated lifespan, whereas treatment with diflunisal resulted in the extension of the lifespan of only V30M treated flies (Fig. 1B). This lifespan extension was not observed in TTR-A flies after treatment with diflunisal (Fig. 1B). Moreover, the remarkable extension of lifespan observed in V30M flies was not accompanied by a significant decrease of TTR deposition or a notable improvement in locomotor abilities when compared to diflunisal-treated TTR-A flies (Fig. 3 and 5). These findings suggest that this increase of lifespan may result from a TTR-independent effect of diflunisal. A previous study on the effect of non-steroidal anti-inflammatory drugs on *Drosophila* found an extension of lifespan and a delay in the age-dependent decline of locomotor activity (Danilov et al., 2015). In light of that study, the evaluation of diflunisal for its capacity to ameliorate locomotor impairment and to expand lifespan in flies should be interpreted with caution.

Our studies aim to expand the therapeutic landscape for neuropathic ATTR. Both liver transplantation and protein stabilization by tafamidis, current lines of treatment for FAP, have shown positive outcomes overall but are limited to very specific patient profiles and present important caveats. Liver transplantation is an invasive procedure only for hereditary ATTR that is not recommended to all patients because of risks associated with the procedure itself, such as organ rejection, or post-surgery complications (Carvalho et al., 2015). For instance, of clinical importance for ATTR patients with cardiac pathology, liver transplantation sometimes fails to stop disease progression and leads to cardiac wild-type TTR deposition and heart failure (Liepnieks and Benson, 2007). TTR stabilization by tafamidis has been approved for the treatment of only FAP-V30M at an early stage because it has proven to be insufficient to stop disease progression at late stages or in patients with cardiac involvement (Bulawa et al., 2012; Lozeron et al., 2013; Fujita et al., 2017). It is imperative to develop additional therapeutic tools for those cases for which stabilization of TTR or liver transplantation may not be fully effective or not recommended.

Inhibition of TTR aggregation and amyloid seeding by peptide inhibitors may represent a novel strategy for halting progression of ATTR, especially for those cases where liver transplantation or stabilization by small compounds may not be sufficient. The peptide inhibitor TabFH1 was designed against the two amyloid-driving segments of TTR to target amyloid growth of existing TTR fibrils (Saelices et al., 2015). The optimization of this original inhibitor led to TabFH2, which inhibits amyloid seeding of wild-type and mutant TTR catalyzed by amyloid fibrils extracted from ATTR patients (Saelices et al., 2018). In the present study, we found that TabFH2 delays ATTR progression in *Drosophila* models whilst decreasing TTR deposition (Fig. 2, 3, 4 and 5). Our initial *in vivo* studies in flies show promising results and encourage continuing the evaluation of peptide inhibitors for the treatment of ATTR in more complex animal models.

## Author contributions

L.S. was responsible for project conception, designed the research, and wrote the manuscript. L.S, M.P. and D.S.E analyzed the data. M.P. and K.P. performed the research. All authors gave final approval of this manuscript.

## Acknowledgments

We thank Manuel Saelices for his help on the computational analysis of 2D trace drawings, and Chi-Hong Tseng for his help on the statistical analysis. We are grateful for support from the Amyloidosis Foundation (Grant #20170827) and NIH (Grant #R01-AG048120).

